# 𝒟-BLUP: a differentiable genomic BLUP model with learnable variance and marker weights

**DOI:** 10.1101/2025.11.25.690542

**Authors:** Joseph Guhlin, Peter Dearden

## Abstract

Genomic best linear unbiased prediction (GBLUP) is widely used for genomic selection in livestock and crop breeding. There is growing interest in connecting machine learning with genomic breeding value prediction. Although the BLUP formula is itself mathematically differentiable, existing implementations do not expose a differentiable computational graph and cannot be trained end-to-end. Here, we present 𝒟-BLUP, a differentiable implementation of the genomic mixed model in JAX that makes the BLUP solve part of the training process rather than an external step. The variance ratio λ and optional block-level kernel weights are treated as trainable parameters, allowing these components to be learned directly from prediction error while preserving the BLUP structure used in animal and plant breeding. This means BLUP can now fit naturally within gradient-based models common in machine learning without losing its interpretability. The method solves the familiar system (*G*_*w*_ + λ*I*) û = *y*\* with either an unweighted VanRaden genomic relationship matrix *G* or a block-weighted variant *G*_*w*_, via automatic differentiation, allowing simultaneous training. On a public dataset, 𝒟-BLUP with a fixed VanRaden kernel reproduces rrBLUP estimated breeding values and predictive performance, while learning λ, and optionally SNP block weights, from a mean squared error objective. 𝒟-BLUP preserves the structure of BLUP used in routine breeding programs but makes it differentiable, maintaining its interpretability while making it embeddable in machine learning pipelines.

## 2. Introduction

### 2.1. Genomic prediction and BLUP

Genomic selection [1] is common in many primary production systems, especially breeding, where genomic information is used to predict breeding values, thus accelerating response to selection [1]. In this setting, linear mixed models, in particular best linear unbiased prediction [2] (BLUP) and its genomic extension [3] (GBLUP), provide the foundational framework for combining fixed effects with genomic random effects in a single model.

The standard mixed model for a single trait can be written as

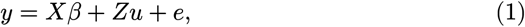

where *y* is the *n* dimensional phenotype vector, *X* is the design matrix for fixed effects with associated coefficients *β, Z* is the design matrix for random effects *u*, and *e* contains residuals. In genomic prediction, the random effects are assumed to follow

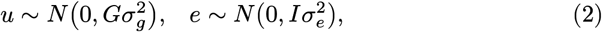

where *G* is a genomic relationship matrix (GRM), 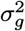 is the additive genetic variance captured by the markers, and 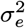 is the residual variance.

In typical GBLUP applications, *G* is constructed from biallelic single-nucleotide polymorphism (SNP) markers using the VanRaden formulation [3], which yields a dense *n* × *n* kernel that captures realized genomic relationships among individuals. The mixed model in Equation 1 and Equation 2 can then be solved either in marker or individual space, and the resulting mixed model equations are well established in breeding practice [2].

### 2.2. GBLUP

Despite its widespread use, GBLUP introduces several practical constraints when used as a component within larger machine learning systems. First, hyperparameter tuning is external to the main optimization loop; the variance components 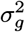 and 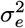, or equivalently the variance ratio 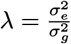, are typically estimated by restricted maximum likelihood (REML) [4], [5] or by cross-validated grid search via specialized software, creating an outer optimization loop that complicates end-to-end training when combined with other differentiable components.

Second, the kernel is typically rigid. The VanRaden GRM assumes that all markers contribute equally after centering and scaling, which is efficient and robust but limits the incorporation of trait-specific weighting or functional priors. Implementations [6], [7] supporting flexible kernels often deviate from the BLUP formulation practitioners are familiar with, or require custom code for each new kernel.

Third, many implementations form *G* explicitly and rely on dense factorizations such as Cholesky decompositions [8]. This restricts scaling as *n* increases, posing challenges for deployment on constrained-memory accelerators (GPUs) and preventing the use of automatic differentiation frameworks. As a result, GBLUP is typically treated as an external tool rather than a native machine learning layer.

### 2.3. Differentiable BLUP

We denote the differentiable BLUP model 𝒟-BLUP (pronounced D-BLUP). Though BLUP is itself differentiable, as a mathematical function of λ and *G*, current software implementations do not expose a differentiable computational graph, preventing gradients from flowing into variance components and kernel parameters. Autodiff frameworks such as JAX [9] allow gradients to flow through the BLUP solve as part of the computational graph, making it straightforward to parameterize λ and marker or block weights, enforce positivity with smooth transforms such as softplus, define prediction-driven losses such as MSE, and backpropagate through the solve step.

The central idea in 𝒟-BLUP is to preserve the mixed model in Equation 1 and Equation 2, including the familiar BLUP system that is solved in individual space, but to express the solve itself as part of a differentiable computation graph. Differentiability enables end-to-end learning of λ and kernel weights, allowing BLUP components to participate directly in gradient-based optimization. The genomic layer then maps phenotypes *y*\* to random effects û by solving

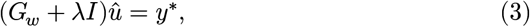

where *G*_*w*_ is either the standard VanRaden GRM *G* or a weighted variant, and λ and the weights are trainable parameters. This turns GBLUP into a differentiable layer that can be embedded in machine learning models and potentially combined with learned embeddings or multiple kernels, and optimized using gradient-based methods.

### 2.4. Contributions of this paper

This paper makes four methodological and implementation contributions.

First, 𝒟-BLUP is a JAX-based implementation of the single-trait GBLUP system that learns the variance ratio λ via gradient-based optimization rather than via grid search or REML. Instead of optimizing λ directly, the model optimizes an unconstrained parameter λ_log_, which is transformed to a strictly positive variance ratio using a softplus function with a small offset for numerical stability (*ε*). JAX automatic differentiation propagates gradients through the linear solve, enabling end-to-end learning of λ.

Second, we extend this to configurable-sized marker blocks (`diff_vr`), a weighted VanRaden kernel, where blocks of markers share non-negative weights learned jointly with λ. This provides a simple mechanism to allocate more or less influence to different genomic regions while retaining the mixed model structure.

Third, 𝒟-BLUP and `diff_vr` are validated on public datasets [10], [11]. For a fixed VanRaden kernel, 𝒟-BLUP reproduces rrBLUP breeding values and predictive performance while learning λ end-to-end. In the weighted case, `diff_vr` learns non-uniform block weights while remaining numerically stable.

Fourth, the implementation is designed for integration into machine learning models. It supports both dense solves for moderate sample sizes and matrix-free conjugate-gradient routines for larger problems, along with preliminary stochastic Lanczos quadrature[12] components for future log-determinant approximations. This provides a validated foundation for scaling to settings where forming a dense *n* × *n* kernel is infeasible, and the design remains compatible with accelerator hardware.

## 3. Methods

### 3.1. Model and notation

For a single trait, let *y* denote the phenotype vector, *X* the design matrix for fixed effects, and *u* the vector of genomic random effects for each individual. The working model is

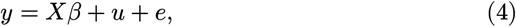

where *β* is the vector of fixed effect coefficients and *e* is a vector of residuals. The random effects and residuals are assumed to follow

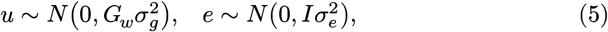

where *G*_*w*_ is either the unweighted VanRaden GRM *G* or a weighted variant. The variance ratio is defined as

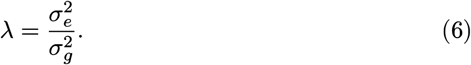

We pre-adjust for fixed effects using OLS, and define

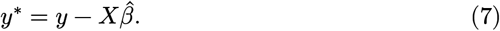

The genomic layer is then fitted to *y*\* with *Z* = *I*. In the dual (“kernel ridge”) formulation, we solve for coefficients *α*:

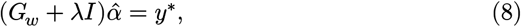

and obtain the BLUP of genomic effects as

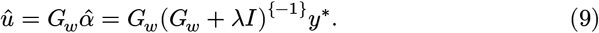

Here *û* are the estimated genomic random effects (estimated breeding values). The same formulation applies to both the fixed-kernel 𝒟-BLUP model and the weighted `diff_vr` model.

### 3.2. VanRaden genomic relationship matrix

The genotype matrix is denoted by *X*, where *X*_*i*,*j*_ counts the number of reference alleles (0, 1, or 2) for individual *i* and SNP *j*. Let *p*_*j*_ denote the reference allele frequency at SNP *j* in the sample.

The genotypes are first standardized to obtain *Z*,

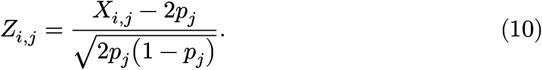

This scaling rescales each SNP to approximately unit variance under Hardy– Weinberg equilibrium and removes the mean.

The VanRaden GRM is then defined as

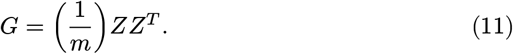

The resulting *G* captures realized genomic relationships among individuals. In the fixed kernel version of 𝒟-BLUP, *G*_*w*_ in Equation 8 is set equal to *G*.

### 3.3. rrBLUP baseline

We use rrBLUP [13] as a reference BLUP implementation. rrBLUP solves the same BLUP system in Equation 8, for a fixed *G* constructed by the VanRaden method and an externally tuned λ.

In rrBLUP, λ is selected via grid search over candidates defined on the log scale. For each λ, rrBLUP solves the system, and we evaluate predictive performance via cross-validation. The value of λ that minimizes validation mean squared error is selected and used to compute the final û_rr_ on the full training set.

This configuration provides a baseline for both predictive accuracy and estimated breeding values. In particular, the correlation between û_rr_ and the 𝒟-BLUP estimates û_𝒟_ is used as a measure of agreement between the differentiable and typical implementations.

### 3.4. 𝒟-BLUP with fixed VanRaden kernel

In the fixed kernel configuration, 𝒟-BLUP uses the VanRaden GRM *G* from Equation 11 as *G*_*w*_ in the BLUP system Equation 8 but treats the variance ratio λ as a trainable parameter.

We parameterize λ using a real-valued unconstrained parameter λ_log_. The variance ratio is obtained by applying a softplus transform and adding a small positive constant *ε* for numerical stability,

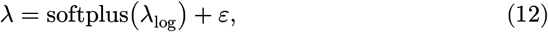

with *ε* typically chosen to be a small scalar such as 10^−6^. This parameterization enforces λ > 0 while allowing unconstrained optimization of λ_log_.

Once λ is set, the BLUP system is solved either directly or iteratively, depending on the dataset size. For the moderate *n* considered in the baseline experiments, a direct Cholesky factorization is sufficient and yields numerically stable solutions. The solution û represents the genomic random effects. Because we train on residual phenotypes *y*\*, the model’s prediction in this space is

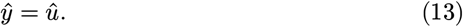

The training loss for 𝒟-BLUP with a fixed kernel is the mean squared error on the training set. This loss is fully differentiable with respect to λ_log_ because both the softplus transform and the linear solve are differentiable operations in JAX.

Gradients of ℒ_MSE_ with respect to λ_log_ are obtained using JAX automatic differentiation. The parameter λ_log_ is updated by Adam [14] or a related first-order optimizer, as described in Section 3.6. The result is an end-to-end training loop that updates λ solely based on predictive performance, without an explicit outer hyperparameter search.

### 3.5. Weighted VanRaden kernel (diff_vr, block mode)

The `diff_vr` variant extends 𝒟-BLUP by introducing block-specific weights that modulate the contribution of subsets of markers to the GRM. This allows the model to allocate higher or lower importance to different genomic regions while preserving the BLUP structure.

Let the *m* SNPs be partitioned into *B* disjoint blocks. For each block *b*, a non-negative weight *w*_*b*_ is defined via a softplus transform,

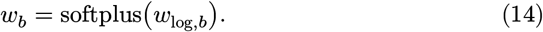

The parameter vector *w*_log_ is unconstrained and trainable. The square root of the block weight is used to rescale the standardized genotype matrix. For each SNP *j* belonging to block *b*, the corresponding column of *Z* is multiplied by 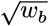 to construct a weighted matrix *Z*_*w*_.

The weighted GRM is then defined as

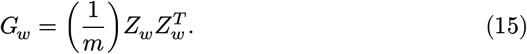

Substituting Equation 15 into Equation 8 yields the standard BLUP system with a weighted kernel. Both λ and the block weights *w*_*b*_ are trained jointly.

The data fit term in the loss is MSE, now with *G*_*w*_ replacing *G*. To prevent unbounded growth or collapse of weights, a quadratic regularisation term is added that encourages weights to remain near 1,

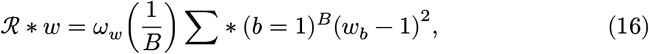

where *ω*_*w*_ ≥ 0 is a regularisation strength. The total training loss is the sum of the MSE term and this regularizer. When *ω*_*w*_ is large, the weights remain close to 1, and `diff_vr` behaves similarly to the fixed kernel 𝒟-BLUP. When *ω*_*w*_ is moderate, the model can deviate from the standard VanRaden kernel to emphasize or de-emphasize particular blocks.

The implementation also supports per-SNP weights with the same parameterization as Equation 14, but in this manuscript, only block-level weights are used to reduce the number of free parameters and facilitate interpretation, as this mode remains experimental.

### 3.6. Training and implementation details

𝒟-BLUP, as well as `diff_vr`, are implemented in JAX and built as Equinox[15] modules so that they behave as standard neural network layers, while optimizers from optax[16] are used for training.

Training uses full-batch updates with gradients computed over the entire training set. This matches the standard mixed-model formulation and avoids the additional variance introduced by mini-batching. The parameters λ_log_ and *w*_log_ are initialized to give a moderate variance ratio and nearly uniform weights, respectively. Concretely, the initial λ in Equation 12 is chosen by a simple variance/REML-style heuristic on the training data, so it is of the same order as standard mixed-model choices, and 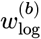 is initialized so that all blocks start from a common baseline with *w*_*b*_ ≈ 1.

We use the Adam [14] optimizer with a fixed learning rate in all experiments. We then train for a fixed number of epochs. For the cross-validation experiments described in Section 5.1, model parameters are reinitialized for each training split to avoid information leakage between folds.

For the moderate sample sizes in our primary experiments, a dense Cholesky factorization is used to solve Equation 8 and the weighted block `diff_vr` model. Our implementation also provides a matrix-free conjugate gradients solver [17] that requires only matrix-vector products with *G* or *G*_*w*_, and stochastic Lanczos quadrature[12] routines for approximating log determinants.

### 3.7. Evaluation metrics

We use several complementary metrics to compare rrBLUP, 𝒟-BLUP, and 𝒟-BLUP with `diff_vr`.

Predictive accuracy is reported as the Pearson correlation between model predictions and target values on held-out individuals. For PIC traits, the targets are estimated breeding values (EBVs), and we report corr(*ĝ, g*) between predicted and reference EBVs. For BF and other traits without separate EBVs, the targets are phenotypes, and we report corr(*ŷ, y*). This correlation is the primary measure of predictive performance.

For traits with very different scales, we additionally consider the normalized quantity 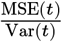 to enable cross-trait comparisons. For methods that produce individual-level random effects, we also report the correlation between their predicted EBVs and the analytic GBLUP solution on the training set as a parity diagnostic, but not as a primary performance metric. For `diff_vr`, we summarize the learned block weights by their mean and range (minimum and maximum) within each cross-validation run, together with the final regularisation parameter λ.

## 4. Data

### 4.1. Datasets

𝒟-BLUP and `diff_vr` are evaluated on two publicly available datasets with individual-level genotypes and phenotypes; one dataset provides five phenotypes, EBVs, and genotypes, referred to as the PIC dataset [10]; from the other, we use a single phenotype, backfat thickness, with covariates from the large white pig data [11].

After standard quality control, the dataset consists of *n*_real_ individuals and *m*_real_ SNP markers. Quality control filters include a minor allele frequency threshold, individual and marker call rate thresholds, and the removal of markers with excessive missingness. Standardization to *Z* follows Equation 10.

The fixed effect design matrix *X* includes an intercept and a set of covariates relevant to the trait and species. These may include sex, herd or flock, birth year or season, and additional management or batch effects, depending on the dataset. Fixed effects are fitted by ordinary least squares, and residuals *y*\* are constructed according to Equation 7.

## 7 Experiments and results

### 5.1. Experimental setup

Our analyses use a 3-fold cross-validation design to compare rrBLUP, 𝒟-BLUP, and diff_vr.

The data are partitioned into *K* approximately equal-sized folds. For each repeat of the cross-validation procedure, one fold is held out as the test set and the remaining *K* − 1 folds form the training set. The same splits are used for all methods to ensure a fair comparison.

For each training split, the rrBLUP baseline is fitted by constructing the VanRaden GRM *G* from the entire training set, performing a grid search over λ and solving Equation 8 at the chosen λ. Predicted phenotypes in the test set are obtained from the resulting û_rr_ and 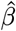.

𝒟-BLUP with a fixed kernel uses the same *G* and *y*\*, but treats λ as a trainable parameter. For each split, λ_log_ is initialized and updated according to the training procedure in Section 3.6. Predictions for the test set are computed using Equation 13.

The `diff_vr` variant uses the same training splits but replaces *G* with the weighted kernel *G*_*w*_ in Equation 15. Block definitions are shared across methods, and `diff_vr` is trained jointly over λ_log_ and *w*_log_ with regularisation strength *ω*_*w*_ in Equation 16. We use block sizes of 500 markers, which are then weighted to generate a GRM.

Predictions, estimated breeding values, and learned hyperparameters (λ and block weights) are stored for each split and method. Evaluation metrics from Section 3.7 are computed on the test sets and summarized across folds (Table 1).

**Table 1.**
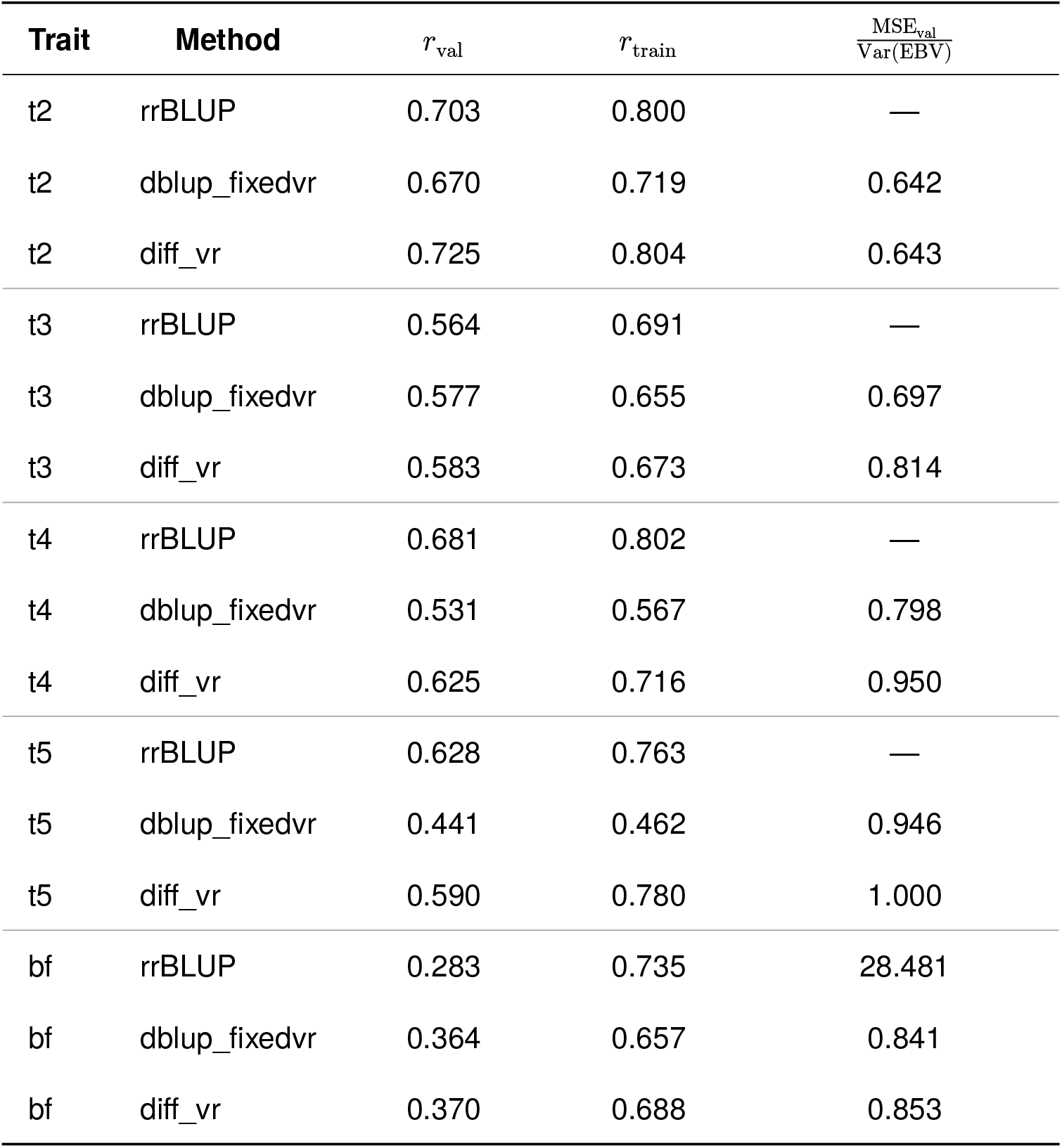
Comparison of Methods across Traits.

### 5.2. 𝒟-BLUP + MLP

For the Large White pig backfat (BF) phenotype, we also evaluate a hybrid model that pairs a genomic BLUP component with a nonlinear covariate term via a small multilayer perceptron (MLP), which maps individual-level covariates to a scalar effect that is jointly summed with the output of 𝒟-BLUP (Figure 1). The BLUP head operates on residuals obtained by subtracting the MLP predictions from the training phenotypes. It then solves the mixed-model system on the training individuals and projects the resulting genomic solution to all individuals via the GRM. The final predictor for each sample is the sum of the covariate MLP output and the genomic BLUP component.

**Figure 1.**
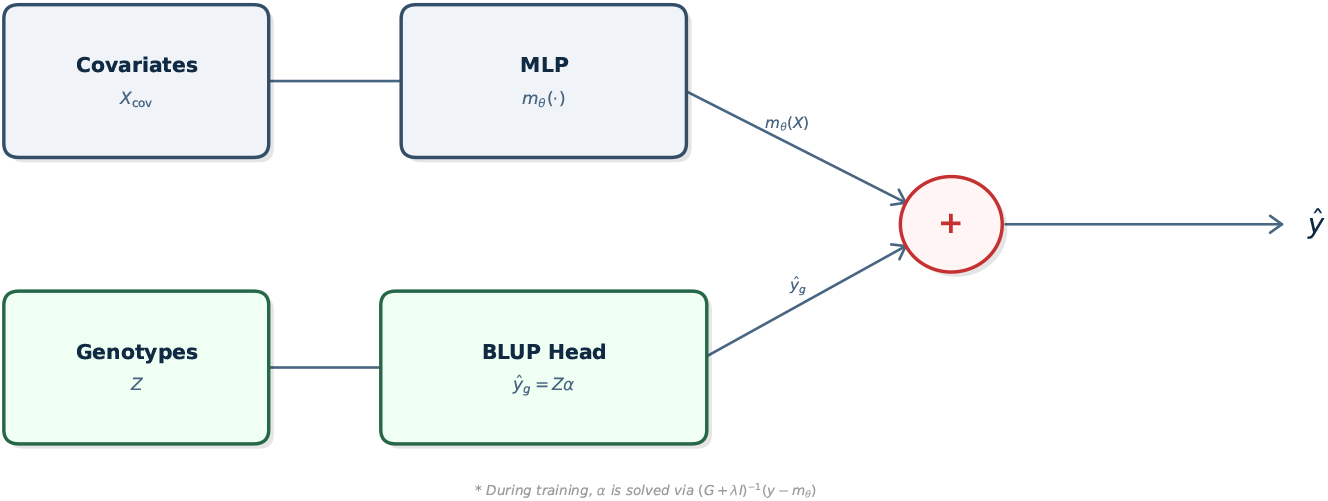
The D-BLUP–MLP architecture consists of two parallel pathways. A covariate pathway maps individual-level covariates through a small multilayer perceptron (*m*_*θ*_). A genomic pathway computes a VanRaden GRM and solves the mixed-model equations on the training residuals 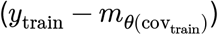. The resulting solution (*α*) is projected to all individuals to obtain genomic predictions 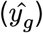. The final predictor combines the covariate and genomic components as 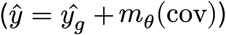. This preserves the additive mixed-model structure while allowing a flexible covariate model in a fully differentiable pipeline.

**Figure 2.**
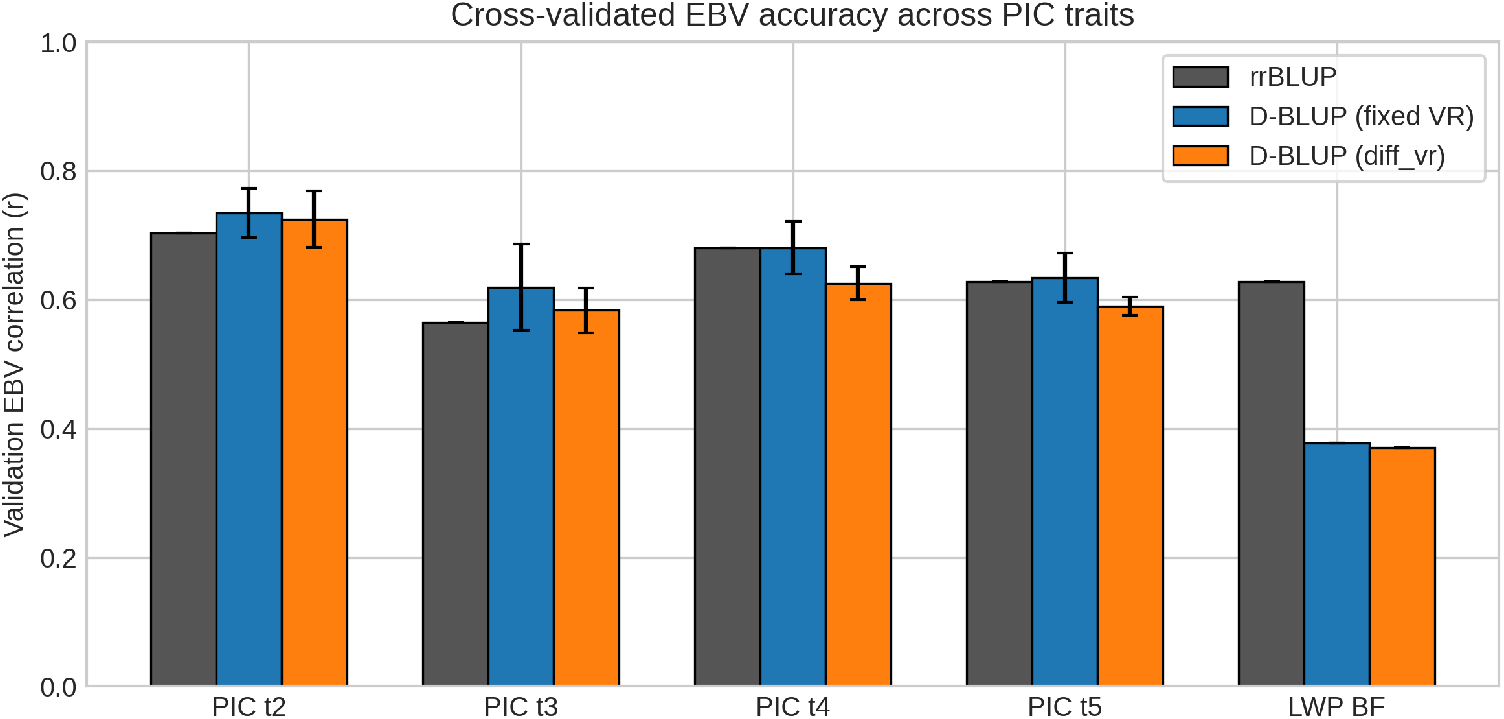
Cross-validated prediction accuracy for rrBLUP, D-BLUP with a fixed VanRaden kernel, and the weighted D-BLUP (`diff_vr`) model across five PIC pig traits. Bars show the mean Pearson correlation between predicted and observed EBVs in the held-out samples, and error bars denote the standard deviation across three cross-validation folds. Across all traits, differentiable BLUP variants perform similarly to rrBLUP, demonstrating parity between the classical and differentiable implementations. The backfat EBV correlation loses accuracy due to a lack of covariate handling in 𝒟-BLUP, but is included here as it is used in Section 5.2.

This retains the linear mixed-model structure while adding a shallow nonlinear covariate term. Both the variance-ratio parameter and the MLP weights are optimized end-to-end against the prediction loss.

On the Large White Pig backfat trait, the BLUP-MLP experiment demonstrates that our differentiable GBLUP layer can be trained end-to-end with a small neural network and yields competitive accuracy. We feed the recorded management/sex covariates through a two-layer MLP, add the resulting nonlinear fixed-effect term to a standard VanRaden GBLUP head, and jointly optimize the MLP weights and the genetic regularisation parameter by gradient descent on phenotypic MSE. With the same corrected phenotypes and covariates, rrBLUP attains a validation correlation of roughly 0.28, a pure dblup_fixedvr GBLUP baseline reaches about 0.36. The MLP+BLUP model reaches ≈0.37–0.38 at its best checkpoint, while keeping λ in a sensible range. This shows that a differentiable BLUP layer alongside a small neural covariate model can exploit nonlinear covariate structure without destabilizing the genetic solver. Combining 𝒟-BLUP with an MLP serves as a proof-of-concept that a BLUP layer and a traditional neural network can be trained simultaneously.

### 5.3. Parity on real data

On the PIC dataset, 𝒟-BLUP, with a fixed VanRaden kernel, matches rrBLUP in both predictive performance and estimated breeding values. Trait t1 is excluded due to very low heritability and low performance with both methods.

Across three cross-validation folds, predictive correlations between Ŷ and *y* in the test set are very similar for rrBLUP and 𝒟-BLUP. Differences in 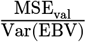 between methods are small relative to fold-to-fold variability and show no consistent direction. This indicates that learning λ from prediction error does not degrade performance relative to grid search.

The learned λ values for 𝒟-BLUP lie within the range of values considered by the rrBLUP grid search and tend to concentrate near the values that minimize validation MSE for rrBLUP (Figure 3). This suggests that gradient-based λ optimization in 𝒟-BLUP recovers the same operating regime as the more expensive grid search.

**Figure 3.**
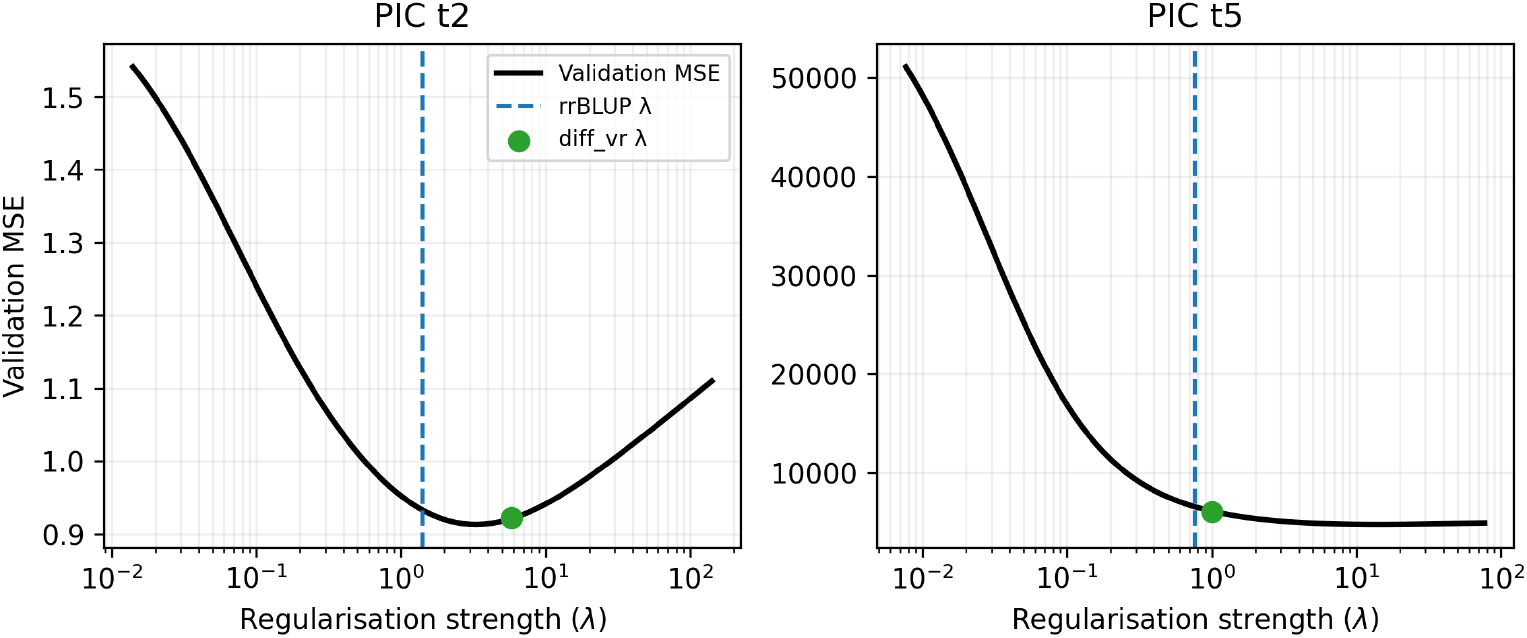
Validation-set mean squared error (MSE) as a function of the regularisation parameter λ on a log_10_ scale for two representative PIC traits. The dashed blue line indicates the λ selected by rrBLUP grid search, and the green point marks the λ learned end-to-end by the `diff_vr` model. The flatness of the loss landscape near the optimum explains why the learned λ values fall within the same region as the grid-search optimum. This illustrates that gradient-based tuning of λ is stable and recovers the same operating regime as rrBLUP.

### 5.4. Behavior of diff_vr

With moderate weight regularisation (intermediate *ω*_*w*_ in Equation 16), `diff_vr` achieves predictive correlations and MSE ratios comparable to those of the fixed-kernel 𝒟-BLUP. The correlation between breeding values from `diff_vr` and from the fixed kernel 𝒟-BLUP remains high, indicating that block weighting does not destabilize the BLUP solutions under this regularisation regime.

Under strong weight regularisation (large *ω*_*w*_), `diff_vr` weights are forced close to one, and the model effectively collapses back to the fixed kernel 𝒟-BLUP. In this regime, predictive performance and breeding values are indistinguishable from those of the unweighted model within numerical tolerance. This shows that the weighted kernel extension can be configured to behave conservatively when prior information about genomic heterogeneity is weak.

## 6. Discussion

𝒟-BLUP is a differentiable implementation of the standard GBLUP model that maintains the linear mixed model structure familiar to professional breeders while exposing variance ratios and marker weights as trainable parameters for machine learning applications. In a public dataset, 𝒟-BLUP with a fixed VanRaden kernel reproduces rrBLUP breeding values and predictive performance to a degree that is negligible for practical decision-making. The variance ratio λ can be learned directly from the prediction error, eliminating the need for an external grid search.

The `diff_vr` extension demonstrates that a weighted VanRaden kernel can be integrated into this framework and trained stably. When weight regularisation is strong, `diff_vr` collapses back to the standard kernel, a safe default when the data do not support meaningful departures from the unweighted VanRaden structure.

For practitioners, 𝒟-BLUP keeps the mixed model formulation while enabling automatic hyperparameter learning and integration with other machine learning components, as an interpretable layer rather than an opaque machine learning system. Future developments will connect the differentiable BLUP layer with richer representations, including sequence-derived embeddings or multi-kernel models, trained with the same gradient-based tools used here. This merges genomic prediction with machine learning without discarding the statistical properties and foundation of BLUP.

This work shows that BLUP can be made differentiable while preserving near-parity with traditional models on single-trait, single-effect problems. More complex genetic architectures, including multiple random effects, genotype by environment interactions, and non-additive effects, will be required to make this usable for breeders. We also show that end-to-end training is stable both for block-weighted kernels and when BLUP is combined with a neural covariate module.

At the same time, the implementation provides a concrete path towards addressing several of these limitations. Matrix-free solvers (CG and SLQ) enable the application of 𝒟-BLUP to populations where the genomic relationship matrix is too large to be explicitly formed. Approximate marginal likelihood objectives based on log-determinant approximations can be incorporated to perform variance-component learning beyond simple MSE-driven tuning. The modular design of the 𝒟-BLUP layer should allow extensions to additional kernels and to multi-trait models used in routine genetic evaluation.

Future work will focus on turning 𝒟-BLUP from a single-trait demonstration into a practical tool for modern breeding datasets. A natural next step is to move beyond simple prediction loss and incorporate likelihood-based objectives, enabling the learning of variance components in richer genetic models without manual tuning. The framework is also well-suited to incorporating alternative genomic kernels and multi-trait structures, allowing the differentiable layer to mirror the models used in routine genetic evaluation. Finally, we intend to apply 𝒟-BLUP to full breeding program data in livestock and crops. Here, matrix-free solvers and distributed computation will matter less as theoretical machinery and more as the practical means to model real-world populations at the scale breeders now work with. We aim to develop a differentiable mixed-model layer that is useful not only as a machine-learning component but as an applied tool for understanding and predicting genetic merit in high-throughput breeding programs.

### 6.1. Conclusion

𝒟-BLUP is a differentiable implementation of the genomic mixed model that learns variance ratios and block-level kernel weights through prediction-driven training. By validating on real livestock data, we show that it matches rrBLUP in performance while enabling integration into gradient-based pipelines.

This work introduces a flexible foundation for hybrid models to combine the interpretability of BLUP with the adaptability of modern machine learning. As the scale of breeding data grows, differentiable solvers and kernel-weighted extensions like `diff_vr` offer a path toward fully trainable genomic prediction tools that remain biologically grounded.

